# Segmenting accelerometer data from daily life with unsupervised machine learning

**DOI:** 10.1101/263046

**Authors:** Dafne van Kuppevelt, Joe Heywood, Mark Hamer, Séverine Sabia, Emla Fitzsimons, Vincent van Hees

## Abstract

**Purpose:** Accelerometers are increasingly used to obtain valuable descriptors of physical activity for health research. The cut-points approach to segment accelerometer data is widely used in physical activity research but requires resource expensive calibration studies and does not make it easy to explore the information that can be gained for a variety of raw data metrics. To address these limitations, we present a data-driven approach for segmenting and clustering the accelerometer data using unsupervised machine learning.

**Methods:** The data used came from five hundred fourteen-year-old participants from the Millennium cohort study who wore an accelerometer (GENEActiv) on their wrist on one weekday and one weekend day. A Hidden Semi-Markov Model (HSMM), configured to identify a maximum of ten behavioral states from five second averaged acceleration with and without addition of x, y, and z-angles, was used for segmenting and clustering of the data. A cut-points approach was used as comparison.

**Results:** Time spent in behavioral states with or without angle metrics constituted eight and five principal components to reach 95% explained variance, respectively; in comparison four components were identified with the cut-points approach. In the HSMM with acceleration and angle as input, the distributions for acceleration in the states showed similar groupings as the cut-points categories, while more variety was seen in the distribution of angles.

**Conclusion:** Our unsupervised classification approach learns a construct of human behavior based on the data it observes, without the need for resource expensive calibration studies, has the ability to combine multiple data metrics, and offers a higher dimensional description of physical behavior. States are interpretable from the distributions of observations and by their duration.

## 1 Introduction

Accelerometers are increasingly used for studying daily physical activity. A common technique to process accelerometer data is the so called ‘cut-points’ approach. This approach allows calculation of time spent with the acceleration registered by the accelerometer between certain thresholds to define physical activity intensity levels (sedentary, light, moderate, vigorous), at different bouts duration [1]. The threshold values used in this approach are calibrated relative to an indirect calorimetry derived metabolic equivalent (MET) level which, in brief, is a proxy for energy expenditure relative to rest [2,3]. Results from the cut-points approach are easy to report and reproduce. However, this approach comes with several challenges. Firstly, a known challenge is the complex relationships of acceleration with energy expenditure, activity types, study populations, and study designs, make that cut-points easily overfit to the experimental conditions under which they are derived. Secondly, the approach involves many parameters, such as bout length, that are often chosen without a clear exercise physiological motivation. Thirdly, the cut-points approach leads to collinearity between classes which partly result from the compositional nature of the data [4] and partly from causal relations between behaviors [5]. This collinearity complicates the study of interactions between behavioral categories [6].

The cut-point approach traditionally used the magnitude of acceleration as its input. The orientation of the accelerometer under static conditions has also emerged as an additional informative metric to detect human posture [7], more recently for the detection of sleep [8], and in the Sedentary Sphere method for detecting sitting behavior and visualizing the data [9]. Further, the data can also be explored using automated methods such as machine learning. Machine learning methods that use labelled data, referred to as supervised machine learning, have previously been used for activity type classification and energy expenditure estimation [10–13]. Although such methods have shown potential for physical activity intensity assessment, they have disadvantages similar to the cut-points approach in that the trained classifier may overfit to the specific experimental conditions under which it was trained. Unsupervised machine learning on the other hand has received less attention in relation to physical activity intensity assessment. These methods are data-driven, allow identification of the characteristic states in the data, and can be applied to free-living data directly. Note that they are called states rather than categories, because they are defined by a Markov model rather than by absolute thresholds. As a result, they do not require time consuming and expensive calibration studies including a year of work to plan and conduct the study, they do not require costs related to exercise laboratory usage, and they may avoid arbitrary decisions in the design of the cut-point approach.

We hypothesize that if such a data-driven approach to segment the data is only provided with input data that has a known physiological meaning like the magnitude of acceleration, it may be possible to learn physiologically meaningful segments in the data. If successful, this would overcome limitations of the methods requiring population sample calibration, such as the cut-point approach. Our primary aim is to implement HSMM to identify states characterized by the intensity of the activity undertaken. There is currently no gold standard method for categorising activity intensity. We therefore assess the comparability of our new approach with the traditional cut-points approach, by looking at collinearity between time spent across different activity intensity states and between cut-point categories. Large collinearity between intensity states or categories complicates modeling behavioral interactions. We will investigate this collinearity with correlation analysis and principal component analysis [6,14–16]. Further, we examined the plausibility of the relation between resulting HSMM-defined activity intensity states, cut-points categories and time-use diary records. A supplementary aim was to assess the influence of adding accelerometer orientation metrics. In this study we used data from 500 fourteen-year-old participants.

## 2 Methods

### 2.1 Participants

Members of the Millennium Cohort Study (MCS), a longitudinal study managed by the Centre for Longitudinal Studies at the UCL Institute of Education participated. The MCS follows over 19,000 young people in the UK born in 2000/1, conducting home interviews every 2 to 4 years. Its sixth survey, at age 14, included the collection of physical activity data using wrist-worn activity monitors [17–19]. The study obtained ethical approval from the National Research Ethics Service (NRES) Research Ethics Committee (REC) – London Central (ref: 13/LO/1786). Written consent was first required from a parent/guardian, and verbal consent from the cohort member [17,19]. A random subset of 500 participants was selected for the present analysis. Millenium Cohort Sixth Sweep data, protocols and metadata are available to the scientific community, under DOI 10.5255/UKDA-SN-8156-3.

### 2.2 Study protocol

Interviewers placed tri-axial accelerometers (GENEActiv) with respondents during home visits and requested them to wear the device on their non-dominant wrist for two complete days; one during the week and one at the weekend, randomly selected at time of placement. Each day lasted for 24 hours: from 4 am in the morning to 4am the following morning. Participants received text messages reminding them to complete the tasks on the selected days. Upon completion of the second day of activity data collection respondents were required to return the accelerometer in a pre-paid envelope. The accelerometer sample frequency was set at 40 Hertz and the dynamic range was ±8*g*. The orientation of the acceleration axes, seen from the anatomical position, is as follows: the x-axis points in medio-lateral direction (direction of thumb), the y-axis in longitudinal direction (direction of middle finger), and the z-axis in dorsal-ventral direction (perpendicular to skin). For further details on the study design and protocol see [18,19]. Additionally, participants were asked to record their categories of behavior in a time use diary[17]. Participants were asked to provide a full record of what they did on the two days (activities), from 4am to 4am the next day, as well as where they were, who they were with, and how much they liked each activity – using pre-coded lists. Participants were offered the choice between an app and online version (with a paper version available for those unable or unwilling to use one of the other two).

### 2.3 Accelerometer data pre-processing

The raw data from the accelerometer is processed with R package GGIR [20] which extracts the two days on which the accelerometer was supposed to be worn (days defined from 4am to 4am). Next, it estimates calibration error based on static periods in the data and corrected if necessary [21], and estimates accelerometer non wear time using a previously reported algorithm [22,23]. The main metrics extracted from the data are five second average of the *vector magnitude of body acceleration* and the *orientation of each of the three acceleration sensors relative to the horizontal plane.* The vector magnitude of acceleration is calculated as Euclidian Norm Minus One (ENMO), which in formula corresponds to 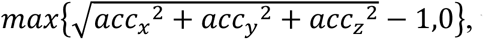 with acc_x_, acc_y_, and acc_z_ referring to the three orthogonal acceleration axes pointing in the lateral, distal, and ventral directions, respectively. From here on, we will refer to the ENMO value as simply *acceleration.* The orientation angles of the acceleration axis relative to the horizontal plane are calculated as
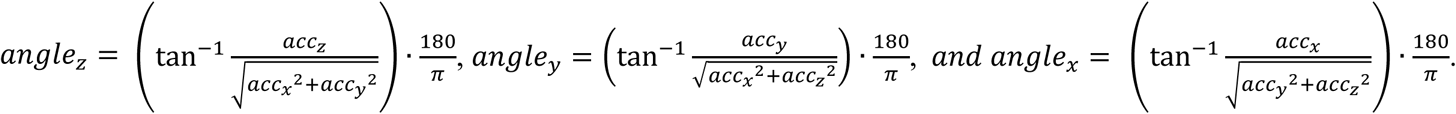

For all continuous time periods with no z-angle change of more than 5 degrees lasting at least 5 minutes, the acceleration values were set to zero to take out the possible influence of increased calibration error during sustained inactivity periods, e.g. as a result of temperature [21]. To account for variation in sign of the signal as a result of wearing the accelerometer upside down, the angles were corrected as follows: If the value for the *x*-*angle* has a positive median during all time periods detected as active (calculation described in the next section) then the device is considered to be worn incorrectly, in that case, the *x*-*angle* and *y*-*angle* (flipped around zero) are negated to mirror the orientation.

### 2.4 Conventional cut-points approach

The acceleration magnitude and angle-z metric are used to assign each of the 5-second epochs to one of the following ten exclusive categories:

- Sustained inactivity: all continuous time periods with no z-angle change of more than 5 degrees lasting at least 5 minutes [8];
- Inactivity: defined, outside identified sustained inactivity, as acceleration below 40 mg, divided into periods of inactivity lasting less than 10 minutes, bouts of inactivity lasting between 10 and 29.9 minutes, and bouts of inactivity for at least 30 minutes;
- Light physical activity (LPA): defined as acceleration between 40 and 120 mg, subdivided into spontaneous LPA lasting less than 1 minute, bouts LPA lasting between 1 and 9.9 minutes, and bouts LPA lasting at least 10 minutes;
- Moderate or vigorous physical activity (MVPA): defined as acceleration above 120 mg, subdivided into spontaneous MVPA lasting less than 1 minute, bouts MVPA lasting between 1 and 9.9 minutes, and bouts MVPA lasting at least 10 minutes.

In order to account for natural variations into acceleration values inside one activity bout, bouts were calculated so that short interruptions accumulate to no more than 20% of the bout length (MVPA) and no more than 10% (light and inactivity bouts). Further, interruptions in bouts are not allowed to last longer than a minute, and interruptions in bouts are counted towards the time spent in bouts over a day. The duration of all bouts was defined as the time elapsed between the start (first epoch meeting the threshold criteria) and the end (last epoch meeting the threshold criteria) of the episode. The bouts were computed with function g.getbout from R package GGIR, metric 4.

In the absence of validated cut-points for our population, we used the reported acceleration values across activity types in children and adults from a study by Hildebrand and colleagues [24]. The choice for three intensity levels, inactivity, light and MVPA, is widely used in the physical activity research community. The sub-classification of these levels in bout durations, was guided by the common practice to look for bouts of at least 10 minute of MVPA [1], and the common practice of looking for bouts of at least 30 minutes inactivity or sedentary behavior [25]. The sustained inactivity category has been shown to be a proxy for sleeping time in adults [8], but could generally be interpreted as time segments without movement and rotation.

### 2.5 Hidden semi-Markov models

The goal of using an unsupervised method is to segment the data in time periods that can be clustered into segments with similar behavior. Hidden Markov Models (HMMs) as used by others for accelerometer data [26,27] do not model the duration of the state. The related Hidden Semi-Markov Models (HSMM) have the advantage of modelling time distribution for the different behavioral states. HSMM have proven to be valuable for similar segmentation tasks in ubiquitous computing [28,29]. In a Hidden semi-Markov model (HSMM) clustering is performed into *hidden states.* The word *hidden* is used because the states are not directly observed, but found by the algorithm. Further, the abstract word *states* is used for the data clusters because we do not know (yet) what physical activity intensity category they represent. However, in practice a state can be interpreted as a physical activity intensity category, with its own characteristic distributions of orientation and acceleration values (the observations that are not hidden) and a characteristic distribution of duration. A graphical representation of the HSMM is visualized in Fig 1.

**Fig 1.**
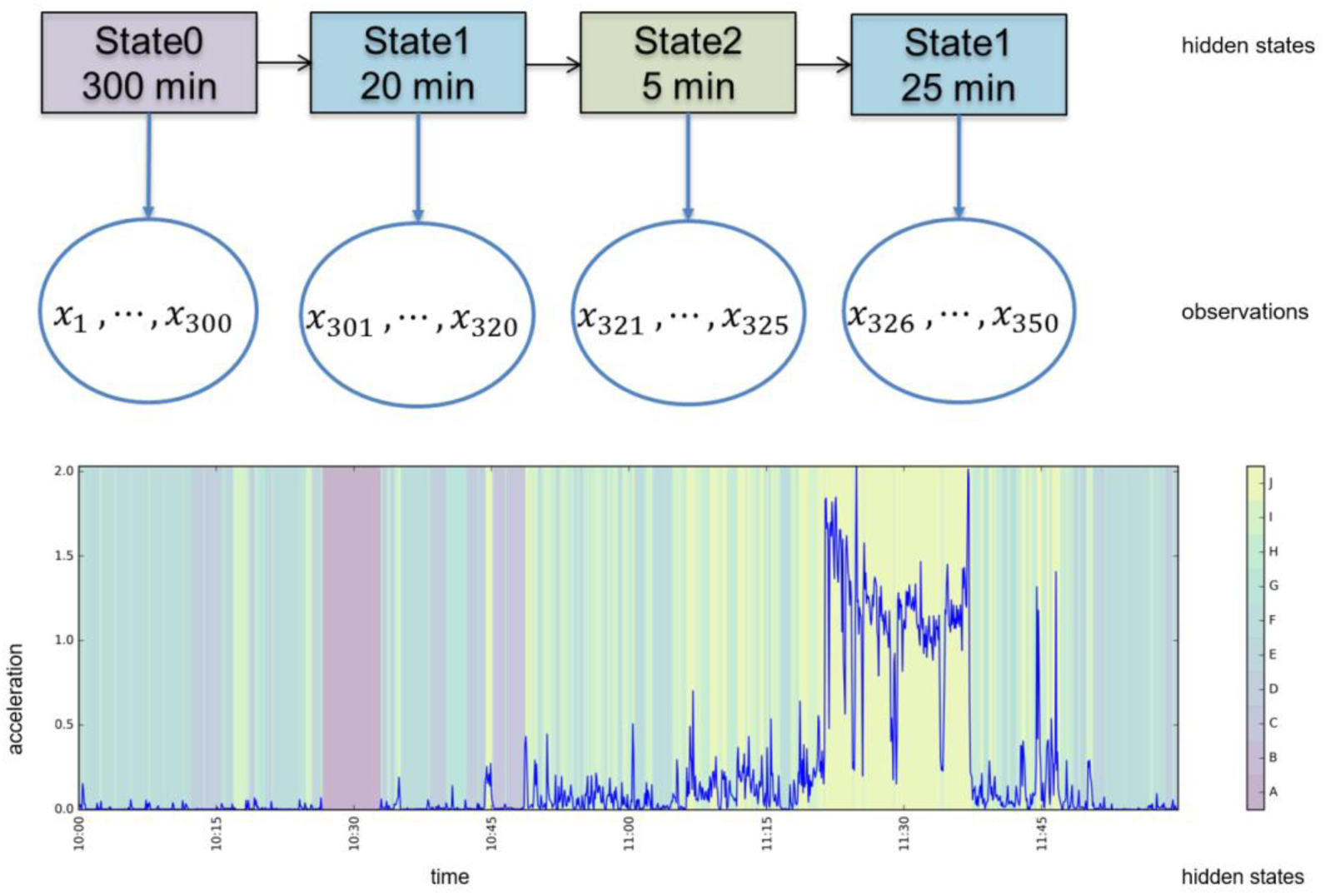
Hidden Semi-Markov model (HSMM), explained schematically (above) and an example of the resulting segmentation (below). The observations (accelerometer metrics), denoted by x, are segmented into states of variable length. Segmentation by the HSMM is based on the model distributions for duration and observations, and transition probabilities between states.

The HSMM is an extension of the widely used Hidden Markov Model [30]. The difference with traditional Hidden Markov models is an explicit distribution for duration of the state. This duration, sometimes called *sojourn time*, is the number of time steps that the model resides in one state before transitioning to the next state.

The observations (acceleration and orientation values) are modelled as Multivariate Gaussian distributions, where each state holds its own mean and variance parameters. The durations are modelled as discrete Poisson distribution, where each state holds its own lambda parameter. In addition, there is a transition probability matrix that indicates how likely it is from each state to transition to each other state.

The parameters of the model (observation distribution parameters, duration distribution parameters, transition matrix), are learned in a Bayesian manner, with a Hierarchical Dirichlet Process HSMM (HDP-HSMM) as presented in [31]. In Bayesian parameter learning, all parameters are represented as prior distributions, and are updated based on the data (Bayesian inference). The specific inference method used for the HDP-HSMM is Gibbs sampling with weak limit approximate algorithm. In Gibbs sampling, each of the parameters is updated iteratively, by sampling from the conditional probabilities for that parameter, based on the current estimated distributions.

In the HDP-HSMM, the transition probabilities between the states are represented as a Dirchlet Process for, in principle, an infinite number of states. However, the number of states is not actually going to be infinite for the following reasons: the Dirichlet Processes share a common parameter that causes the model to favor a small number of states; the weak limit approximate algorithm assumes a maximum number of states (needs to be provided by the user), and the actual number of states is inferred by the model with states being dropped if there are no transitions going in and out of them. Therefore, the number of states can be smaller than the maximum number of states defined by the user.

The forward-backward algorithm calculates the distribution over states, conditioned on the observed data and all model parameters. It serves as a step in the Gibbs sampling, and is used to determine optimal state sequence for a given observation sequence. In the forward pass of this algorithm, the initial state assignment and duration is randomly sampled from the distribution. Note that this random sampling can result in slightly different state assignments in multiple runs of the algorithm.

#### 2.5.1 HSMM parameters

There are several choices that influence the runtime of training the model. Since the forward-backward algorithm is used in each iteration of the training with Gibbs sampling, the complexity of this algorithm contributes to the total training time. The forward-backward algorithm has a runtime of *0*(*T* · *N* · *d_max_* + *T* · *N*^2^) [31]. In this formula, *T* is the length of the sequence, *N* is the number of states and *d_max_* is a maximum chosen duration length. The maximum state duration *d_max_* is a user defined input to control training time. We set *d_max_* at 720 five second epochs, which corresponds to 60 minutes.

The maximum number of states *N_max_* is an input for the weak limit algorithm. Although the number of states is inferred by the algorithm, it can be useful to limit the number of states. It is more computationally efficient to have a small number of states, and easier to interpret the resulting states. *N_max_* = 10 was evaluated, similar to the number of cut-points categories. The number of iterations of Gibbs sampling needs to be chosen so that the algorithm converges to a stable parameter set. Early stopping is introduced when the hamming distance between the assigned states of two consecutive iterations is smaller than 0.05. In other words, convergence is reached if not more than 5% of the time steps are assigned a different state than in the previous iteration. Further, we chose a maximum of 15 iterations.

Lastly, the metrics that are used as observations in the model can be varied. We experimented with two models with different observations as input. The first model used only acceleration, the second model used acceleration together with *angle_x_*, *angle_y_* and *angle_z_*. We will further refer to the resulting models as the *acceleration* model and the *acceleration*+*angles* model, respectively. The information used *acceleration* model is most comparable to the cut-points approach, since the cut-points approach only uses angle for one out of the ten categories. It is not possible in HSMM to instruct the model to use specific variables for only a single state. For the implementation of the Bayesian HSMMs the python package **pyhsmm** (https://github.com/mattjj/pyhsmm) is used, the code is available on Github ([32] and [33]).

### 2.6 Evaluation

In the following sections, we use the term ‘categories’ to distinguish those slices for the cut-points approach and we use the term ‘states’ to discriminate between those slices for the HSMM approach.

Time spent in each state per day and cut-points category was calculated for participants with full 24 hours of data. Principal component analysis was used on the time spent in states and cut-point categories separately. Next, the cumulative variance of principal components, starting at the first principal component, was used to quantify the number of principal components needed to explain at least 95% of the variance across all principal components. This number of required principal components was used as an indicator for the information dimensionality produced by the cut-points approach, HSMM approach with only *acceleration*, and the HSMM approach with *acceleration*+*angles*, in order to assess whether the angle variable added to the dimensionality of the activity pattern. The PCA statistics give an indication of collinearity between the variables: the more components are needed to explain the variance of the data, the less colinear the variables are. As an alternative way of looking at information dimensionality we also looked at the cross correlation between states and between cut-point categories.

Next, to assess comparability of the cut-points and HSMM approaches, correlation coefficients were calculated between time spent in states and cut-points categories, grouped by acceleration level. We expect that the states come with a plausible variation with respect to the conventional method output. To investigate the differences, a descriptive comparison was done of HSMM states, acceleration values, angle values, cut-points categories, and time use diary records. To ease interpretation, only days of measurement with full 24 hours of valid data (no accelerometer non-wear time) were considered for the descriptive comparison [22,34]. We will focus mostly on the HSMM models using *acceleration*+*angles* since we expect the addition of angle variables to give extra insights, *acceleration* only results will be reported in the supplement.

We choose a sample size of 500 participants for two reasons. First, we would like to demonstrate that the HSMM also works in relatively small samples, such that it can be applied in wide range of study conditions not limited to larger cohorts. Cut-points are typically derived from laboratory studies with less than 100 participants by which 500 participants is still a large sample size. Second, adding more data would increase the computing time. To evaluate that a subsample can generalize to a larger population we tested the reproducibility. A HSMM model was trained on a random subset of 250 participants out of the total set of 500, Nmax = 10, using only the *acceleration* variable since this makes it easier to compare state distributions. The states of the original model trained on 500 participants and the model trained on the subset were sorted on acceleration mean so that a match between corresponding states of both models could be made. The model parameters (mean and sigma for the observation distributions, lambda for the duration distributions) of the corresponding states of the two models were compared. Evaluation were based on the closeness of distributions for each state using the Kullback-Leibler divergence, which is an information-theoretic measure to calculate the distance between two statistical distributions [35]. The Kullback-Leibler divergence is denoted with KL(P|Q), where P is the distribution of the observed values as learned by the original model on 500 participants, and Q the corresponding distribution for the subset model.

## 3 Results

A total of 9122 participants accepted to wear the accelerometer, 4970 participants returned the accelerometer and time use diary, out of which a random subsample of data from 500 participants was used for the present study. The demographics of this sample are shown in Table 1. Note that the data of all 500 participants, regardless of whether the data was complete, could be used for training the HSMM model.. The demographics of the participants who wore the accelerometer for 24 hours on both days are also shown in Table 1. In the calculation of the acceleration values on average 31% (standard deviation 5%) of the of the epochs were replaced by zero.

**Table 1:** Characteristics of subsets

|  |  | Data for training | Data for comparisons |
| --- | --- | --- | --- |
| N |  | 500 | 200 |
| Sex |  | 42.4% male | 41.5% male |
| BMI (kg/m<sup>2</sup>) |  | 21.3 ± 4.2 | 21.2 ± 3.9 |
| Country | England | 59.8% | 61.5% |
|  | Northern Ireland | 10.9% | 8.5% |
|  | Scotland | 12.5% | 12.0% |
|  | Wales | 16.8% | 18.0% |
| Equivalized household income quintiles from lowest to highest | 1 | 10.1% | 7.0% |
|  | 2 | 13.1% | 13.5% |
|  | 3 | 20.4% | 21.5% |
|  | 4 | 29.1% | 25.5% |
|  | 5 | 27.3% | 32.5% |
| Valid data per day (hours) |  | 21.0 (IQR: 20.2-24.0) | 24.0 (for all) |
| MVPA guideline met* |  | 38.8% | 44.6% |
| Acceleration (mg): Mean ± standard deviation, |  | 31 ± 13 | 34±13 |

[IQR: Inter quartile range; \* Percentage of participants that spent at least 60 minutes in MVPA on at least one day;
\*\* calculated with metric ENMO, see text]

### 3.1 Correlation analyses and PCA

Ten different states were found by the HSMM training for both the models using *acceleration* only and using *acceleration*+*angles*. The percentage of explained variance for the number of principal components of the derived time use variables is plotted in Fig 2. The lower the curve is, the more dimensions of information the dataset has. Four, five, and eight principle components were needed to explain 95% of the variance of the variables for the cut-points categories (10 variables were input), *acceleration* HSMM model (10 variables were input), and *acceleration*+*angle* HSMM model (10 variables were input), respectively. The absolute correlation between time spent in the 10 different cut-points defined physical activity categories was 0.27±0.24 (range: 0.01-0.85), for the states of the *acceleration* model it was 0.25±0.18 (range: 0.00-0.77) and for the *acceleration*+*angles* model it was 0.20±0.13 (range: 0.01-0.52), as illustrated in Fig 3. For the cut-points categories, there were 8 combinations of variables (out of 45 combinations) that had an absolute correlation of more than 0.5, for the *acceleration* model and the *acceleration*+*angles* model there were 6 and 1 combinations respectively.

**Fig 2.**
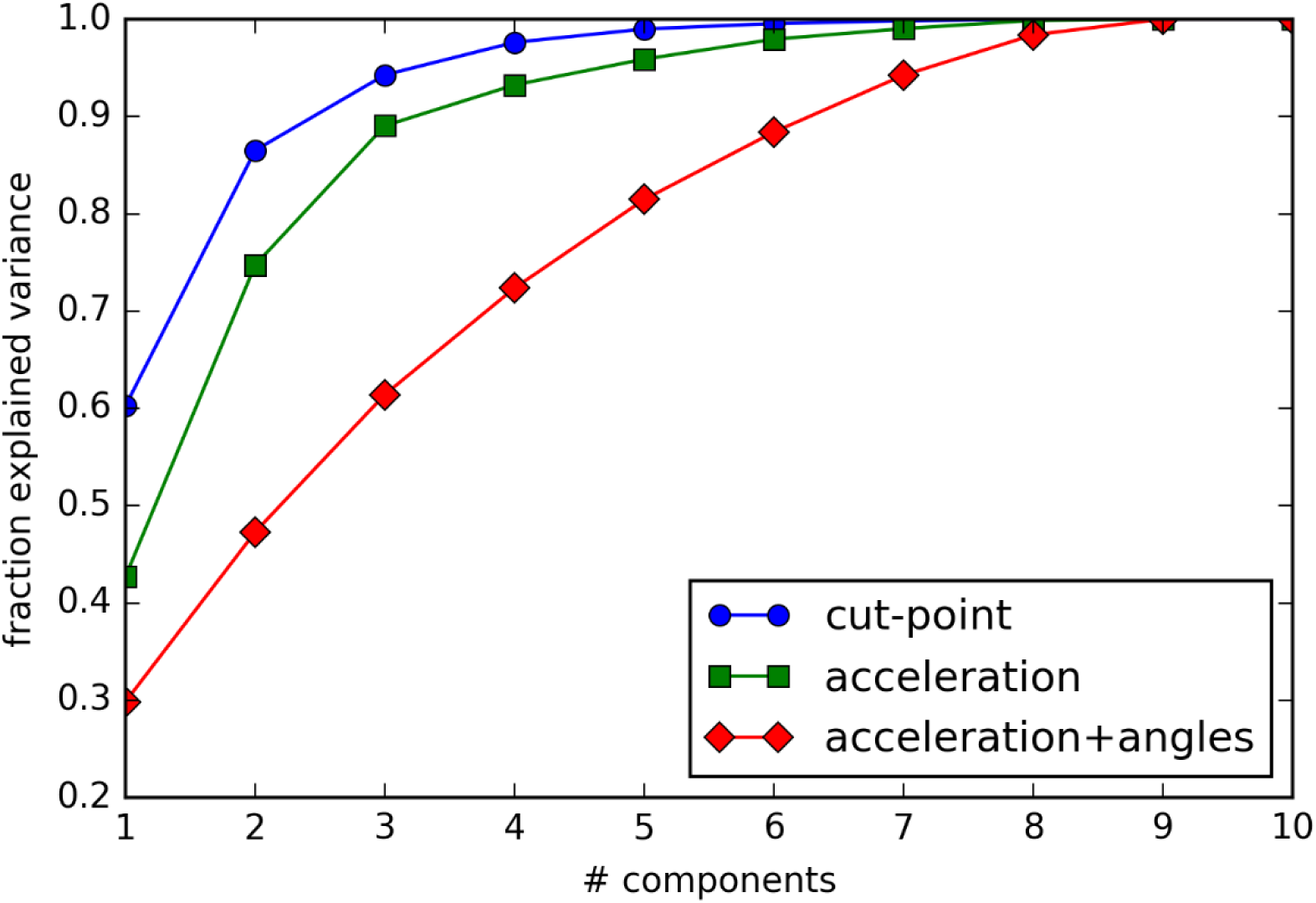
Scree plot showing the explained variance for principal components of time spent in cut-points categories and HSMM states for the HSMM based on acceleration and the HSMM based on acceleration+angles.

**Fig 3.**
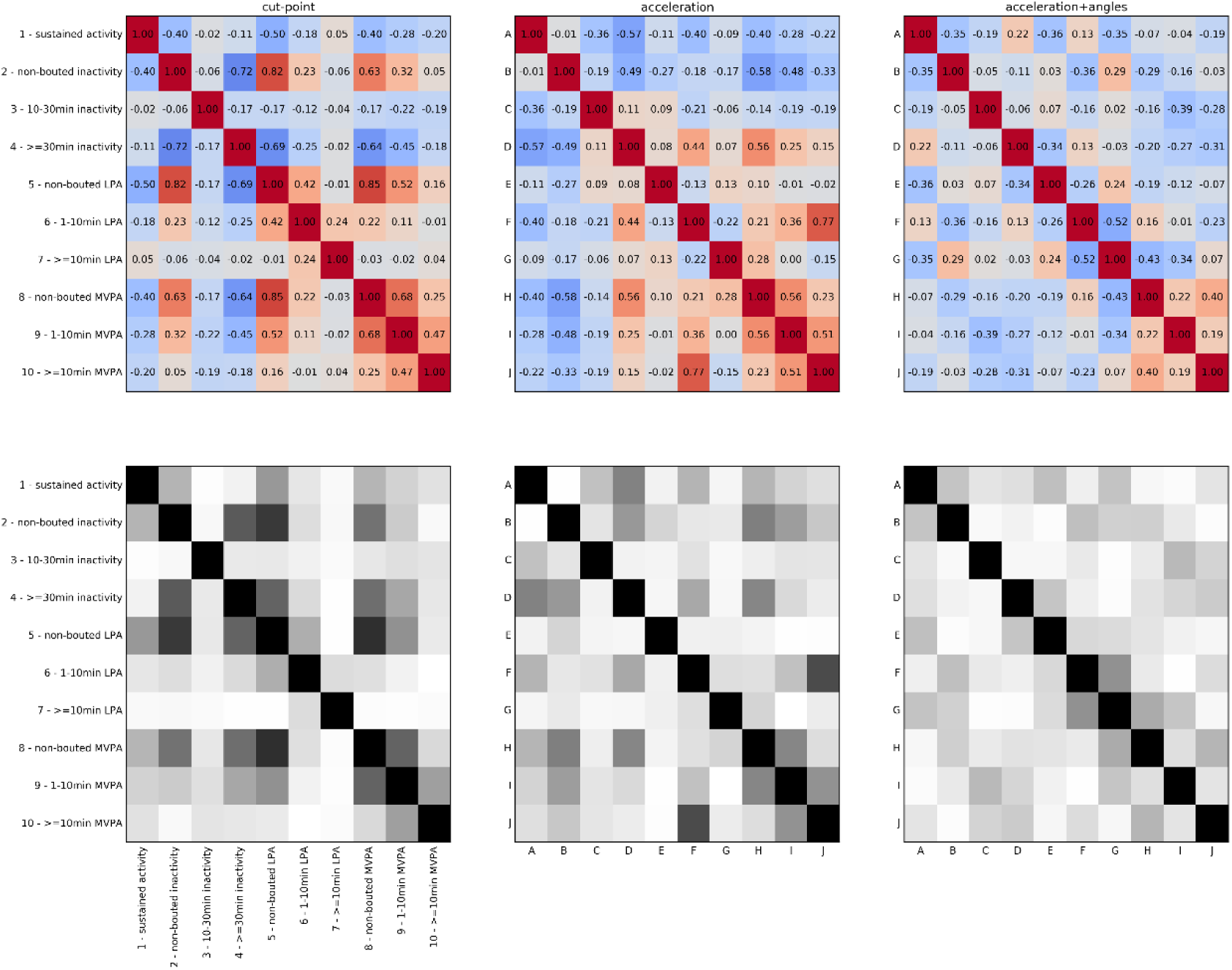
Correlation (top) and absolute value of correlation (bottom) matrices for cut-points categories and model states. The values in the bottom plots are not shown, but can be derived from the top plots by the reader.

### 3.2 Comparison of acceleration and angles to cut-points states

Distributions in acceleration values, angle values, and durations varied by state and threshold categories, see Fig 4. We labeled the states with letters in increasing order of average acceleration. In Table 22 we see how much time the participants spent on average in each cut-points category and each state. A detailed description of the states in the *acceleration*+*angles* model is given in the supplement. We also compare the values in Table 2 for parallels between states and cut-points categories. The same can be done for the *acceleration* model (available in S2 Table of the supplement).

**Fig 4.**
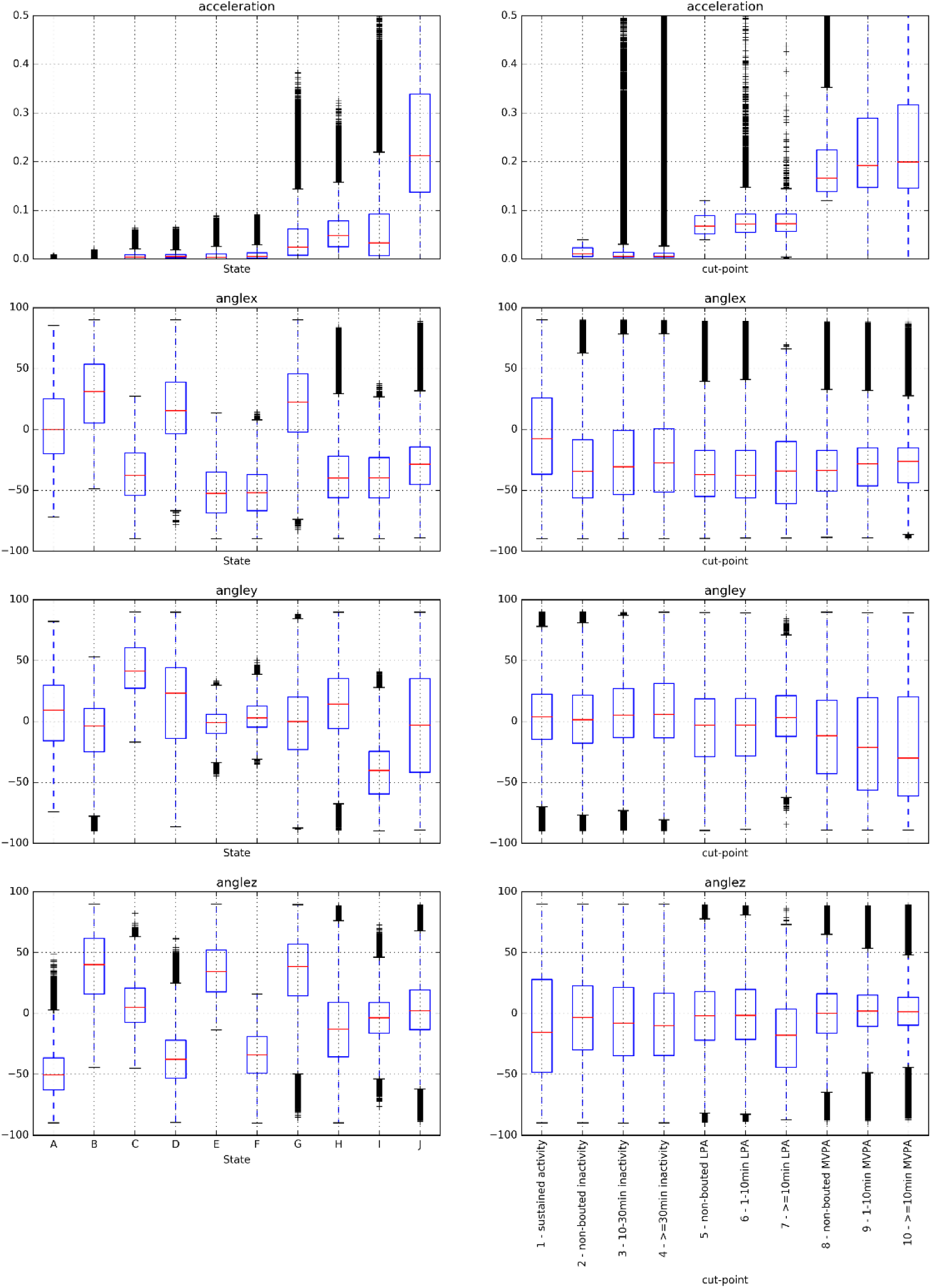
Boxplots for acceleration and angle values per HSMM state (acceleration+angles model, left column) and cut-points category (right column)

**Table 2:** Average time spent (minutes) per participant per day in each state from the acceleration+angle method and cut-points category. The states are sorted on mean acceleration, resulting in higher numbers around the diagonal.

|  |  | state |  |  |  |  |  |  |  |  |  |  | Avg<br>acc |
| --- | --- | --- | --- | --- | --- | --- | --- | --- | --- | --- | --- | --- | --- |
|  |  | A | B | C | D | E | F | G | H | I | J | total |  |
| cut-points category | 1 - sustained inactivity | 197 | 105 | 29 | 0 | 43 | 37 | 2 | 5 | 13 | 0 | 432 | 0.0 |
|  | 2 - non-bouted inactivity | 2 | 6 | 32 | 30 | 44 | 52 | 28 | 41 | 47 | 2 | 284 | 14.5 |
|  | 3 - 10-29.9min inactivity | 3 | 7 | 23 | 26 | 23 | 33 | 12 | 15 | 15 | 1 | 156 | 13.3 |
|  | 4 - >=30min inactivity | 4 | 12 | 46 | 50 | 34 | 59 | 18 | 27 | 28 | 2 | 280 | 13.3 |
|  | 5 - non-bouted LPA | 0 | 0 | 0 | 0 | 2 | 3 | 19 | 66 | 43 | 7 | 142 | 71.6 |
|  | 6 - 1-9.9min LPA | 0 | 0 | 0 | 0 | 0 | 0 | 2 | 6 | 5 | 1 | 15 | 75.1 |
|  | 7 - >=10min LPA | 0 | 0 | 0 | 0 | 0 | 0 | 0 | 0 | 0 | 0 | 0 | 77.3 |
|  | 8 - non-bouted MVPA | 0 | 0 | 0 | 0 | 0 | 0 | 5 | 10 | 19 | 23 | 57 | 220.4 |
|  | 9 - 1-9.9min MVPA | 0 | 0 | 0 | 0 | 0 | 0 | 1 | 3 | 9 | 15 | 28 | 286.7 |
|  | 10 - >=10min MVPA | 0 | 0 | 0 | 0 | 0 | 0 | 1 | 2 | 6 | 9 | 17 | 310.5 |
| total |  | 206 | 131 | 130 | 106 | 146 | 185 | 88 | 175 | 186 | 60 | 1412 |  |
| avg acc |  | 0.0 | 0.6 | 6.5 | 7.4 | 7.8 | 9.6 | 42.5 | 55.9 | 58.2 | 305.2 |  |  |

The model states and cut-points categories can both be grouped in combinations of respectively states and categories that have similar acceleration levels, corresponding to sustained inactivity, inactivity, LPA and MVPA, see Table 2. States were grouped as follows: the group {A, B} corresponds to *sustained inactivity*, states {C, D, E, F} to *inactivity*, states {G, H, I} to *LPA* and state {J} to *MVPA.* The Pearson’s correlation coefficient between the time spent per grouped cut-points category and the time spent per grouped states were 0.69, 0.56, 0.72 and 0.88, respectively for each level of activity defined.

The cut-points categories differed in duration by design (standard deviation of duration means=15.9 minutes), while less difference was observed in durations between states (Fig 5) as there were all shorter than 17 minutes and standard deviation of the duration means was 4.3 minutes for the *acceleration* model and 2.9 minutes for the *acceleration*+*angles* model. In contrast, the distribution of angle values is different for the states with similar acceleration levels. The average values in the cut-points categories for the x, y and z angles have a standard deviation of respectively 8.1, 9.3 and 6.6 degrees over all cut-points categories, whereas over the states this is 31.7, 21.6 and 31.7 degrees.

**Fig 5.**
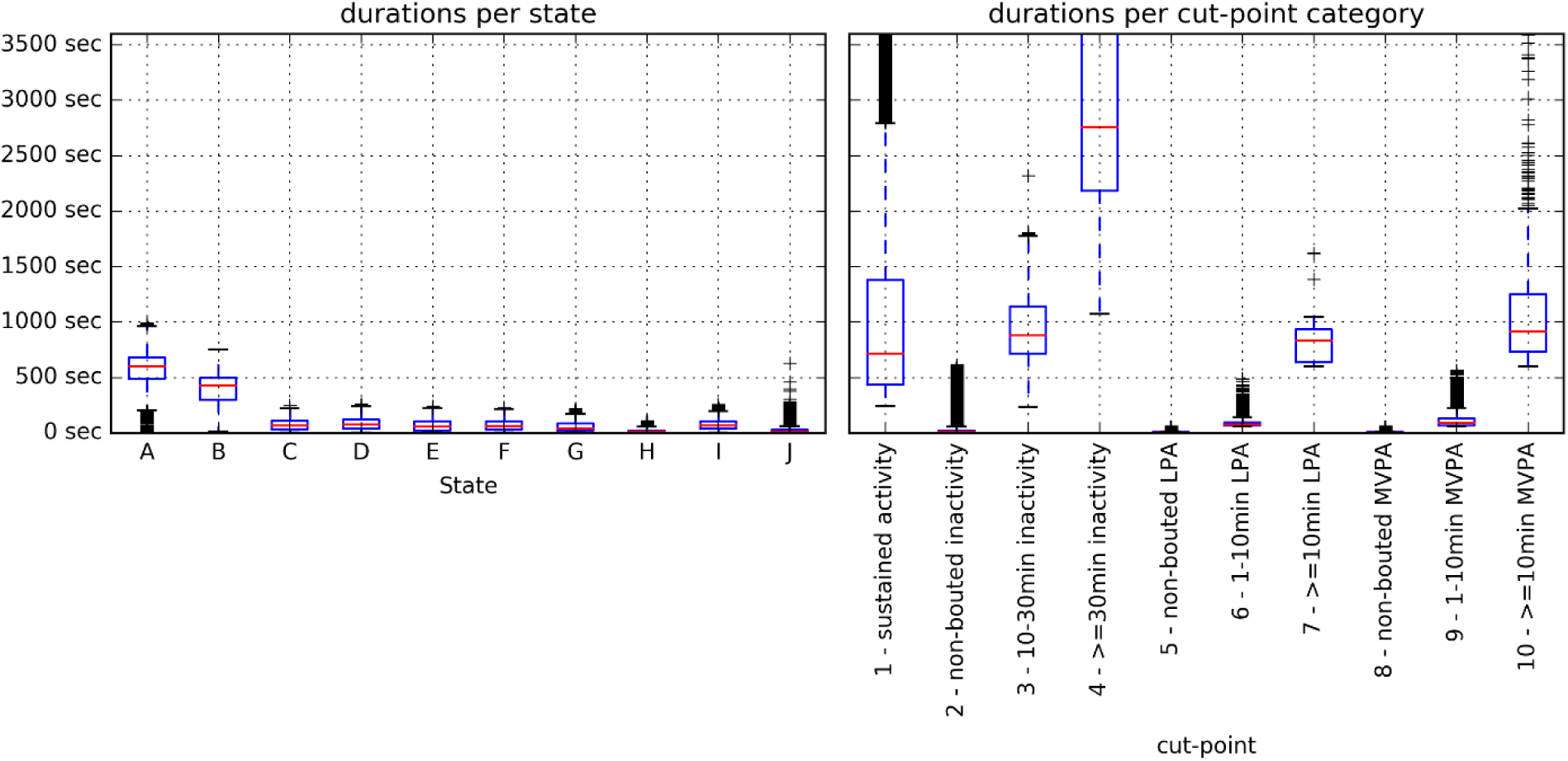
Durations for each HSMM state (acceleration+angles model) (left) and each cut-points category (right)

### 3.3 Comparison against time use diary

To assess whether there is a relation between the states and activities reported in the time use diary, we focused on the 10 most common activities reported in the time use diary, see Table 3 for a comparison with the *acceleration*+*angles* model (the same table for the *acceleration* model is available as S3 Table in the supplement). The sustained inactivity states {A, B} are mostly present during sleep. Strongly sedentary activities, such as “Watching tv ..” and “Playing electronic games..” have most time steps in the inactivate states {C, D, E, F}, which is consistent with our comparisons against the cut-points approach. States {G, H, I} present more time in activities such as speaking or eating a meal. State J appears as a mix of short time in different activities.

**Table 3:** Average time spent (minutes) per participant per day in each state (acceleration+angles model) and the top 10 activities

|  | state |  |  |  |  |  |  |  |  |  |  |
| --- | --- | --- | --- | --- | --- | --- | --- | --- | --- | --- | --- |
|  | A | B | C | D | E | F | G | H | I | J | total |
| **Sleeping and resting (including sick in bed)** | 132.2 | 81.3 | 27.2 | 30.8 | 41.7 | 45.8 | 17.9 | 21.8 | 26.6 | 4.7 | 429.9 |
| **In class** | 0.9 | 1.5 | 8.0 | 6.2 | 10.0 | 20.5 | 8.0 | 19.9 | 18.6 | 6.4 | 99.9 |
| **Watch TV, DVDs, downloaded videos** | 4.4 | 4.4 | 12.7 | 11.3 | 8.8 | 12.8 | 7.2 | 12.6 | 11.7 | 3.4 | 89.4 |
| **Speaking, socialising face-to-face** | 1.5 | 1.0 | 7.7 | 4.1 | 6.8 | 7.0 | 4.0 | 10.5 | 10.4 | 3.6 | 56.4 |
| **Personal care (including taking a shower/bath, grooming, getting dressed etc.)** | 2.9 | 2.7 | 5.2 | 3.6 | 5.3 | 5.8 | 4.3 | 8.3 | 8.4 | 2.9 | 49.4 |
| **Eating a meal** | 1.0 | 1.2 | 5.6 | 4.1 | 5.9 | 6.7 | 4.0 | 8.5 | 8.6 | 3.0 | 48.6 |
| **Playing electronic games and Apps** | 1.7 | 1.4 | 6.6 | 4.6 | 6.3 | 8.4 | 3.4 | 6.8 | 6.7 | 1.7 | 47.6 |
| **Other activities not listed** | 1.2 | 1.0 | 3.0 | 2.0 | 3.5 | 3.7 | 2.8 | 6.0 | 6.8 | 2.3 | 32.2 |
| **Homework** | 1.2 | 1.0 | 3.8 | 2.5 | 3.6 | 4.8 | 2.3 | 4.8 | 4.3 | 1.3 | 29.6 |
| **Travel by car, van (including vehicles owned by friends and family)** | 0.5 | 0.6 | 2.5 | 1.6 | 3.4 | 3.9 | 2.7 | 5.7 | 5.6 | 2.1 | 28.6 |

### 3.4 Reproducibility

The states of the *acceleration* model trained on a subset of 250 participants were sorted on acceleration mean and then matched with the states from the model trained on the full data set of 500 participants. The model parameters (acceleration mean and variance, duration mean and variance) are plotted in Figs 6 and 7. The Kullback-Leibler (KL) divergence for the acceleration distributions is below 1.0 for all state combinations except the two states with small durations. The duration distributions are less consistent over the two models, with 4 out of 10 state combinations having a KL divergence of larger than 1. The calculated KL divergences are listed in the supplement in S4 Table.

**Fig 6.**
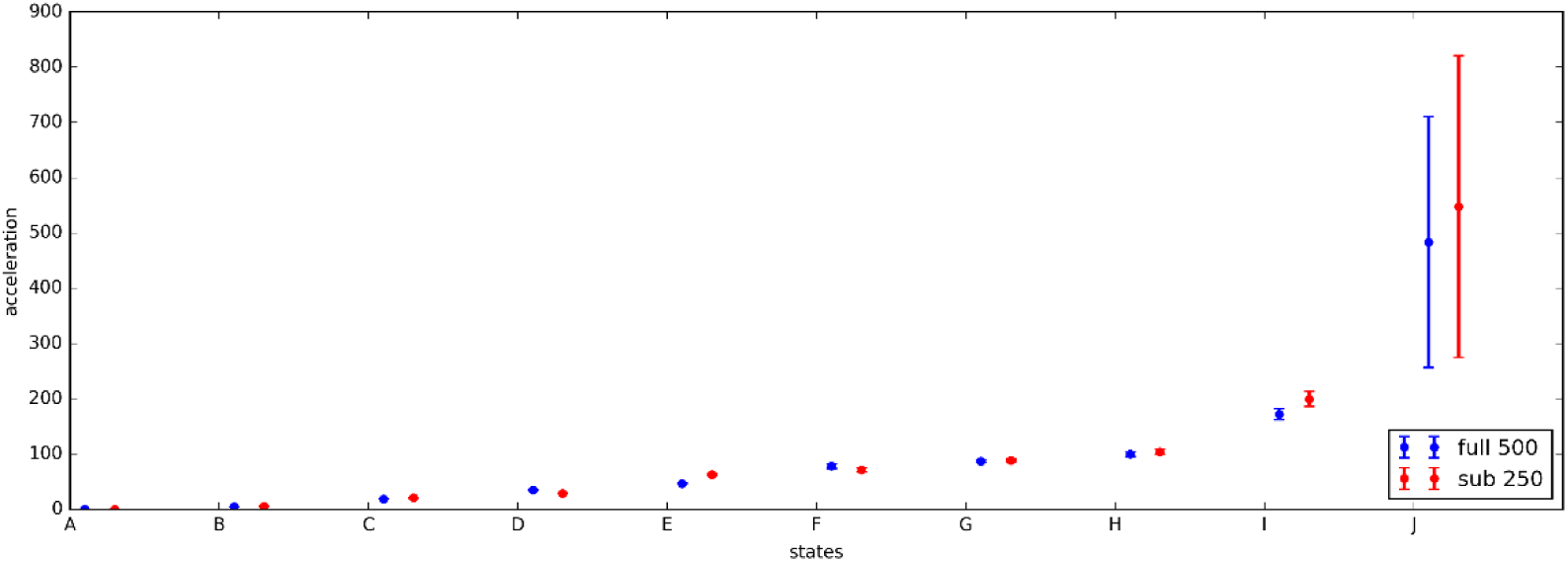
Mean and variance parameters for acceleration.

**Fig 7.**
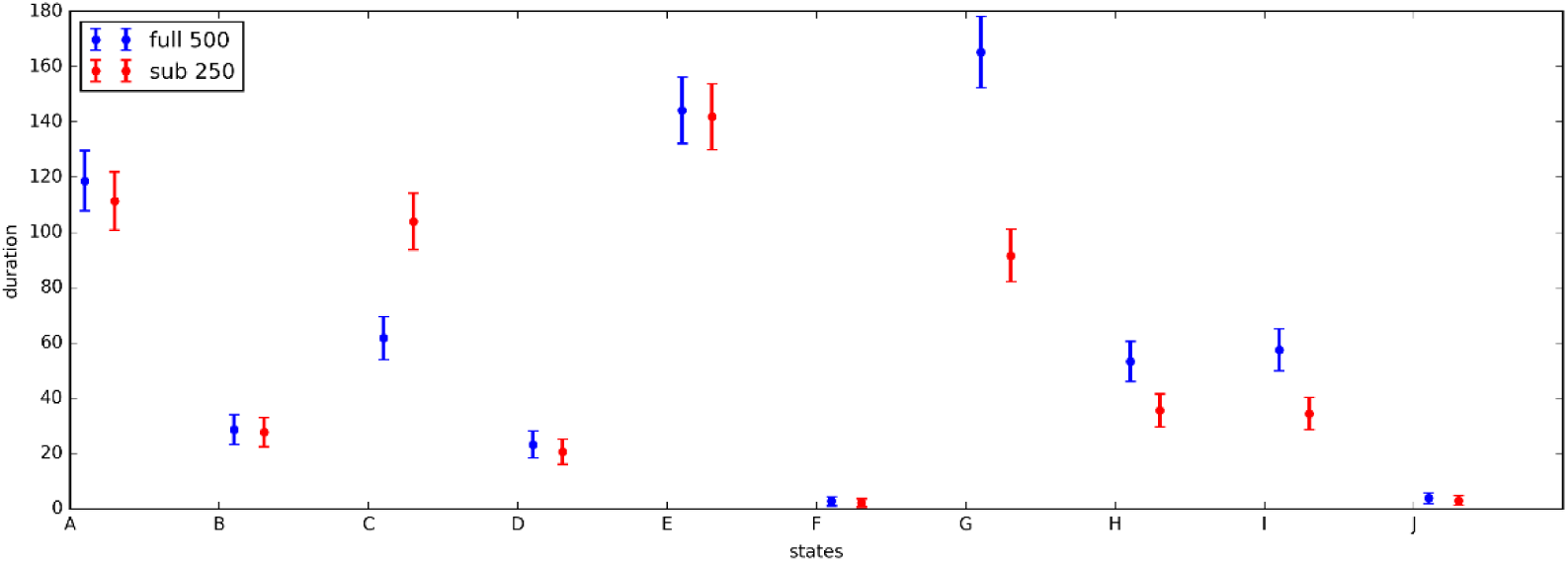
Lambda and variance of parameters of the duration distribution.

## 4 Discussion

This study proposes an unsupervised Hidden semi-Markov Model (HSMM) to segment and cluster data from wearable accelerometers. The HSMM is a possible alternative for the widely-used cut-points approach as it does not require resource expensive calibration studies. The increased use of raw data accelerometry in recent years has resulted also in the need for a new calibration study for every population and every new acceleration processing metric to be used when relying on the cut-points approach [36–41]. Furthermore, these calibration studies are typically limited to small numbers of participants and activity types, thus generalization to different populations and conditions is a known problem [42]. The HSMM approach, examined in the present study, learns ‘states’ from the data, which are described by the mean and variance of the observations (accelerometer derived time series) and by the lambda of their Poisson distributed duration. An important strength of our implementation of the HSMM is that it relies on a small set of input metrics that are relatively easy to interpret in the domain of movement intensity. The HSMM has this advantage of interpretability over other approaches, such as Deep Neural Networks. We intentionally do not use other signal features, because that would introduce the risk that the HSMM model detects activity types rather than intensity. Activity type is a different dimension of physical activity and mistakenly classifying types would undermine interpretation. Therefore, our comparison with the time use diary should be interpreted with care as time use categories do not only reflect variations in intensity.

Our findings show that the HSMM derived states were related to cut-points categories. For example, the HSMM found short lasting states with high acceleration and long lasting states with low acceleration, which is consistent with data derived from cut-points approaches. Further, when states and cut-points categories are grouped by acceleration level, correlations of 0.56 and higher are observed between th e two approaches. The mean acceleration for some of the states was close to the threshold value used in the cut-points approach, which indicates disagreement on the range or distribution of acceleration for typical behaviors. The distributions of durations in the HSMM states is also different from the cut-point categories. However, these differences may well be explained by the fact that the HSMM is driven by the distributions in observations (accelerometer recording) from data collected outside a laboratory in the daily life of British teenagers, while the cut-points approach in our case was driven by cut-points derived from Norwegian children and adults performing a very specific set of activity types in a laboratory setting. By showing most states to last less than 17 minutes, our results contribute to the debate about current practice to quantify physical activity and inactivity in ten minute bouts [43] and thirty minute bouts respectively [44]. Consequently, it may not be surprising that no perfect agreement is found between the cut-points approach and the HSMM approach. More importantly, the HSMM covers a plausible range of acceleration levels (low, medium, and high), durations (from less than a minute to more than 30 minutes), and to some extent, although difficult to interpret, angle ranges. Further, it was reassuring to observe that the principal component analyses applied to time spent in states has a less steep scree plot compared to time use variables based on the cut-points approach. This indicates that research on interactions between behaviors will be less challenged by collinearity. An important strength of the HSMM is that it can account for multi-variate input, even if no prior theory exists on why or how the additional input variables could contribute. In contrast, the cut-points approach needs such a theory [45]. Our PCA results indicates that adding the angles offers a description of physical activity with less collinearity and possibly higher dimensions compared to using acceleration only.

The HSMM approach allows us to move towards a description of physical behavior based on, and driven by, the accelerometer data that can feasibly be collected in both small and large scale populations. The HSMM is not biased by the subjective nature of self-report methods, avoids the complexities of accounting for inter-individual variation in body composition in energy expenditure estimation and the variation in the relationship between body composition and energy expenditure between activity types, and avoids the difficulties with generalizability of supervised learning techniques that rely on training data composed of small numbers of participants and/or activity types. The HSMM approach will not directly fit into the research framework aimed at providing public recommendation on layman constructs like steps or minutes of moderate to vigorous physical activity per day. However, HSMM may speed up and facilitate a data driven approach that could help to understand how variations in activity characteristics, as measured by acceleration and arm angle, relate to health and disease.

### Strengths and limitations

The agreements between time spent in the HSMM states and time use diary categories were poor. Reasons for this are likely to rely on the fact that the time use diary collects broad information on activity type (10 items) and activity context using low time resolution (10-minute slot) in comparison to the physical activity intensity construct as measured by the cut-points approach. Nevertheless, the time use diary allowed us to evaluate how these two constructs relate to one another. A challenge in the design of the present study was that there exists no gold standard for intensity profiling of physical activity in a real life setting in a representative population. Therefore, we combined a variety of analyses to assess the comparability of HSMM and cut-points approach from different perspectives. Future studies might complement the evaluation of models trained on real life data, with data from lab studies, where activities and energy expenditure can be directly measured with indirect calorimetry [24,40,46]. The states found by the HSMM are based on patterns in the data and are not specific for the research question of a user. Therefore, when using the HSMM states as physical behavior descriptors in further research, it might be good practice to undertake post-processing of the data. This includes, for example, grouping states that have similar characteristics regarding the research question (e.g. similar acceleration level), or using the majority state for a larger window to adjust for the desired time granularity.

In practice, it seems not to be feasible to let the model converge to very consistent state assignment (e.g. < 1% Hamming distance). It is not clear from theory how much of this is attributed to variance in the data, and how much could be gained with more training iterations. Therefore, future research (both empirical and theoretical) is needed to investigate the relationship between data size, population characteristics, and convergence.

Future research is needed to better understand the application of HSMMs on physical activity data. For example, the number of states can be varied to optimize face validity, while retaining interpretability and feasibility in terms of training time. The same holds for the size of the input time frame: instead of 5-second time frames, smaller or larger time frames can be used as input. It is also possible to include more input metrics in the model, although that may also complicate the interpretation of the states. In the present study we limited the number of metrics to facilitate a standardized comparison with the cut-points approach and to facilitate interpretation. The use of the z-angle for sustained inactivity detection in the cut-points approach does not undermine the standardized comparison, because the HSMM model also uses this information: When calculating the magnitude of acceleration that is used as input for the HSMM model, values are replaced by zero when the z-angle is constant for a five minutes. The use of different distributions to represent the data in the HSMM model could be investigated, such as a log-normal distribution for the acceleration metric.

The use of metrics that describe the orientation of the device imposes challenges. Firstly, the interpretation of the distributions of the angle values is difficult, asking for visualization and comparison to specific activities to distinguish between states with different angle distributions. Secondly, the correct wear position of the device becomes crucial. In this work, we took a heuristic approach at correcting for improper worn devices. Future research is needed to build a reliable classifier that determines the wear position if signal metrics are used that are body side dependent [47].

Separation of gravitational and calibration of acceleration sensor data is a known challenge [21,22] and the estimates of acceleration are not entirely free from calibration error. By replacing the magnitude of acceleration by zero for the time segments where the accelerometer does not change orientation we ensure that: calibration error as a function of accelerometer orientation does not influence the segmentation of the acceleration data; the contribution of white signal noise to the magnitude of acceleration is minimized, and; bias caused by calibration error is as close to zero across the recording.

We chose a random subset of 500 participants in this study, because we wanted to demonstrate that the HSMM method works in relatively small data sets. Suitability for small datasets is important for uncommon study populations, including the very old, rare diseases, and populations in hard to reach rural areas. The reproducibility experiment suggests that the model for a smaller subset of 250 approaches the model trained on the data of all 500 participants, except for the rarest states. This provides us with confidence that the model generalizes well to more data from the same target group and is not overfitted to the data it is trained on. However, the question remains how much the model generalizes to other populations, e.g. age groups or countries. If enough data are available, it is also possible to train the model for a specific person; this would however make it more difficult to relate the resulting states among the participants. This is a problem of finding the right balance between a model optimized in specific population, and a model that representative for the general population.

The cut-points approach has known limitations, but despite these limitations it has been of tremendous value to the physical activity research community for decades. Therefore, we felt it important to aim for comparability with the traditional approach, while at the same time trying to address one of its limitations. Future analyses to compare the associations of time spent in categories assessed by the cut-point approach and states assessed by the HSMM with health outcomes in different population setting will allow to assess the value of the HSMM approach for research. We conclude that applying Hidden Semi-Markov Models results in informative states, based on the data from a real-world setting. It is possible to relate the states to conventional cut-points categories, to interpret the meaning of the states. The unsupervised model can easily incorporate multiple input metrics, so that the states provide a higher dimensional description of physical behavior.

## 6 List of supplements

S1 Appendix. Supplementary results

S2 Table. Average time spent (minutes) per participant per day in each state from the acceleration method (sorted by mean acceleration) and cut-points category

S3 Table. Average time spent (minutes) per participant per day in each state (acceleration method) and the top 10 activities, acceleration model

S4 Table. Kullback-Leibler divergences for the distributions on the full model and a subset model

S5 Table. Distribution parameters of the HSMM (acceleration) model, for each state

S6 Table. Distribution parameters of the HSMM (acceleration+angles) model for each state

## 5 References

1. Menai M, van Hees VT, Elbaz A, Kivimaki M, Singh-Manoux A, Sabia S. Accelerometer assessed moderate-to-vigorous physical activity and successful ageing: results from the Whitehall II study. Scientific reports. 2017 Apr 3;8:45772.

2. Diaz KM, Krupka DJ, Chang MJ, Kronish IM, Moise N, Goldsmith J, et al. Wrist-based cut-points for moderate- and vigorous-intensity physical activity for the Actical accelerometer in adults. Journal of sports sciences. 2017 Feb 23;1–7.

3. McGarty AM, Penpraze V, Melville CA. Calibration and Cross-Validation of the ActiGraph wGT3X+ Accelerometer for the Estimation of Physical Activity Intensity in Children with Intellectual Disabilities. PloS one. 2016;11(10):e0164928.

4. Chastin SFM, Palarea-Albaladejo J, Dontje ML, Skelton DA. Combined Effects of Time Spent in Physical Activity, Sedentary Behaviors and Sleep on Obesity and Cardio-Metabolic Health Markers: A Novel Compositional Data Analysis Approach. PloS one. 2015;10(10):e0139984.

5. Westerterp KR. Pattern and intensity of physical activity. Nature. 2001 Mar 29;410(6828):539.

6. Guinhouya CB, Soubrier S, Vilhelm C, Ravaux P, Lemdani M, Durocher A, et al. Physical activity and sedentary lifestyle in children as time-limited functions: usefulness of the principal component analysis method. Behavior research methods. 2007 Aug;39(3):682–8.

7. Veltink PH, Bussmann HJ, de Vries W, Martens WJ, Van Lummel RC. Detection of static and dynamic activities using uniaxial\naccelerometers. IEEE Transactions on Rehabilitation Engineering. 1996;4(4):375–85.

8. van Hees VT, Sabia S, Anderson KN, Denton SJ, Oliver J, Catt M, et al. A Novel, Open Access Method to Assess Sleep Duration Using a Wrist-Worn Accelerometer. PloS one. 2015;10(11):e0142533.

9. Rowlands A V, Yates T, Olds TS, Davies M, Khunti K, Edwardson CL. Sedentary Sphere: Wrist-Worn Accelerometer-Brand Independent Posture Classification. Medicine and science in sports and exercise. 2016 Apr;48(4):748–54.

10. Rosenberg D, Godbole S, Ellis K, Lacroix A, Natarajan L, Kerr J. Classifiers for Accelerometer-Measured Behaviors in Older Women. Med Sci Sports Exerc. 2017;49(3):0–0.

11. Ellis K, Kerr J, Godbole S, Lanckriet G, Wing D, Marshall S. A random forest classifier for the prediction of energy expenditure and type of physical activity from wrist and hip accelerometers. Physiological measurement. 2014 Nov;35(11):2191–203.

12. Lyden K, Keadle SK, Staudenmayer J, Freedson PS. A method to estimate free-living active and sedentary behavior from an accelerometer. Medicine and science in sports and exercise. 2014 Feb;46(2):386–97.

13. Freedson PS, Lyden K, Kozey-Keadle S, Staudenmayer J. Evaluation of artificial neural network algorithms for predicting METs and activity type from accelerometer data: validation on an independent sample. Journal of applied physiology (Bethesda, Md: 1985). 2011 Dec;111(6):1804–12.

14. Fung TT, Hu FB, Holmes MD, Rosner BA, Hunter DJ, Colditz GA, et al. Dietary patterns and the risk of postmenopausal breast cancer. International journal of cancer. 2005 Aug 10;116(1):116–21.

15. Lord S, Chastin SFM, McInnes L, Little L, Briggs P, Rochester L. Exploring patterns of daily physical and sedentary behaviour in community-dwelling older adults. Age and ageing. 2011 Mar;40(2):205–10.

16. Varraso R, Fung TT, Barr RG, Hu FB, Willett W, Camargo CA. Prospective study of dietary patterns and chronic obstructive pulmonary disease among US women. The American journal of clinical nutrition. 2007 Aug;86(2):488–95.

17. Ipsos MORI; CLS. Millennium Cohort Study Sixth Sweep (MCS6): Time Use Diary Documentation [Internet]. 2016 [cited 2017 Nov 3]. Available from: http://www.cls.ioe.ac.uk/shared/get-file.ashx?id=3208&itemtype=document

18. Chatzitheochari S, Fisher K, Gilbert E, Calderwood L, Cleary A, Chatzitheochari S, et al. Measuring young people’ s time-use in the UK Millennium Cohort Study: A mixed-mode time diary approach A mixed-mode time diary approach. 2015;

19. Ipsos MORI, CLS. Millennium Cohort Study Sixth Sweep: Technical report (Version2) [Internet]. Vol. 2017. 2017 [cited 2017 Nov 3]. Available from: http://www.cls.ioe.ac.uk/shared/get-file.ashx?id=3284&itemtype=document

20. van Hees V, Fang Z, Zhao JH, Heywood J, Sabia S. R package GGIR. 2017.

21. van Hees VT, Fang Z, Langford J, Assah F, Mohammad A, da Silva ICM, et al. Autocalibration of accelerometer data for free-living physical activity assessment using local gravity and temperature: an evaluation on four continents. Journal of applied physiology (Bethesda, Md: 1985). 2014 Oct 1;117(7):738–44.

22. van Hees VT, Gorzelniak L, Dean León EC, Eder M, Pias M, Taherian S, et al. Separating movement and gravity components in an acceleration signal and implications for the assessment of human daily physical activity. PloS one. 2013 Jan;8(4):e61691.

23. Sabia S, van Hees VT, Shipley MJ, Trenell MI, Hagger-Johnson G, Elbaz A, et al. Association between questionnaire- and accelerometer-assessed physical activity: the role of sociodemographic factors. American journal of epidemiology. 2014 Mar 15;179(6):781–90.

24. Hildebrand M, Van Hees VT, Hansen BH, Ekelund U. Age-Group Comparability of Raw Accelerometer Output from Wrist- and Hip-Worn Monitors. Medicine & Science in Sports & Exercise. 2014 Feb;(accepted for publication 2014):1.

25. Bellettiere J, Winkler EAH, Chastin SFM, Kerr J, Owen N, Dunstan DW, et al. Associations of sitting accumulation patterns with cardio-metabolic risk biomarkers in Australian adults. PloS one. 2017;12(6):e0180119.

26. Lin JF-S, Kulić D. Automatic Human Motion Segmentation and Identification using Feature Guided HMM for Physical Rehabilitation Exercises. Workshop on Robotics for Neurology and Rehabilitation, IEEE International Conference on Intelligent Robots and Systems. 2011;33–6.

27. Pober DM, Staudenmayer J, Raphael C, Freedson PS. Development of novel techniques to classify physical activity mode using accelerometers. Med Sci Sports Exerc. 2006;38(9):1626–34.

28. Duong T V., Bui HH, Phung DQ, Venkatesh S. Activity Recognition and Abnormality Detection with. Proceedings of the 2005 IEEE Computer Society Conference on Computer Vision and Pattern Recognition, IEEE, Washington, D C. 2005;838–45.

29. Van Kasteren TLM, Englebienne G, Kröse BJA. Activity recognition using semi-Markov models on real world smart home datasets. Journal of Ambient Intelligence and Smart Environments. 2010;2(3):311–25.

30. Yu SZ. Hidden semi-Markov models. Artificial Intelligence. 2010;174(2):215–43.

31. Johnson MJ, Willsky AS. Bayesian Nonparametric Hidden Semi-Markov Models. arXiv preprint arXiv:12031365. 2013;14:673–701.

32. van Kuppevelt D, van Hees V. hsmm4acc. 2017.

33. van Hees V, van Kuppevelt D. millenniumcohort-acc. 2017;

34. van Hees VT, Renström F, Wright A, Gradmark A, Catt M, Chen KY, et al. Estimation of daily energy expenditure in pregnant and non-pregnant women using a wrist-worn triaxial accelerometer. PloS one. 2011 Jan;6(7):e22922.

35. Kullback S, Leibler RA. On Information and Sufficiency. The Annals of Mathematical Statistics. 1951 Mar;22(1):79–86.

36. Bakrania K, Yates T, Rowlands A V, Esliger DW, Bunnewell S, Sanders J, et al. Intensity Thresholds on Raw Acceleration Data: Euclidean Norm Minus One (ENMO) and Mean Amplitude Deviation (MAD) Approaches. PloS one. 2016;11(10):e0164045.

37. Vähä-Ypyä H, Vasankari T, Husu P, Mänttäri A, Vuorimaa T, Suni J, et al. Validation of Cut-Points for Evaluating the Intensity of Physical Activity with Accelerometry-Based Mean Amplitude Deviation (MAD). PloS one. 2015;10(8):e0134813.

38. Esliger DW, Rowlands A V, Hurst TL, Catt M, Murray P, Eston RG. Validation of the GENEA Accelerometer. Medicine and science in sports and exercise. 2011 Jun;43(6):1085–93.

39. Hildebrand M, VAN Hees VT, Hansen BH, Ekelund U. Age group comparability of raw accelerometer output from wrist- and hip-worn monitors. Medicine and science in sports and exercise. 2014 Sep;46(9):1816–24.

40. Hildebrand M, Hansen BH, van Hees VT, Ekelund U. Evaluation of raw acceleration sedentary thresholds in children and adults. Scandinavian journal of medicine & science in sports. 2016 Nov 22;

41. Phillips LRS, Parfitt G, Rowlands A V. Calibration of the GENEA accelerometer for assessment of physical activity intensity in children. Journal of science and medicine in sport / Sports Medicine Australia. 2013 Mar;16(2):124–8.

42. van Hees VT, Golubic R, Ekelund U, Brage S. Impact of study design on development and evaluation of an activity-type classifier. Journal of applied physiology (Bethesda, Md: 1985). 2013 Apr;114(8):1042–51.

43. Saint-Maurice PF, Troiano RP, Matthews CE, Kraus WE. Moderate-to-Vigorous Physical Activity and All-Cause Mortality: Do Bouts Matter? Journal of the American Heart Association. 2018 Mar 22;7(6).

44. Kim Y, Welk GJ, Braun SI, Kang M. Extracting objective estimates of sedentary behavior from accelerometer data: measurement considerations for surveillance and research applications. PloS one. 2015;10(2):e0118078.

45. Rowlands A V, Olds TS, Hillsdon M, Pulsford R, Hurst TL, Eston RG, et al. Assessing Sedentary Behavior with the GENEActiv: Introducing the Sedentary Sphere. Medicine and science in sports and exercise. 2013 Nov 20;

46. Brandes M, VAN Hees VT, Hannöver V, Brage S. Estimating energy expenditure from raw accelerometry in three types of locomotion. Medicine and science in sports and exercise. 2012 Nov;44(11):2235–42.

47. Rowlands A V, Olds TS, Bakrania K, Stanley RM, Parfitt G, Eston RG, et al. Accelerometer wear-site detection: When one site does not suit all, all of the time. Journal of science and medicine in sport. 2017 Apr;20(4):368–72.

